# Non-invasive measures of DNA methylation capture molecular aging in wild capuchin monkeys

**DOI:** 10.1101/2025.06.03.657357

**Authors:** B Sadoughi, R Hernández-Rojas, H Hamou, R Lopez, M Mah, E Slikas, SMV Simmons, JD Orkin, JP Higham, SF Brosnan, KM Jack, FA Campos, N Snyder-Mackler, AD Melin

## Abstract

Elucidating the socio-ecological factors that shape patterns of epigenetic modification in long-lived vertebrates is of broad interest to evolutionary biologists, geroscientists, and ecologists. However, aging research in wild populations is limited due to inability to measure cellular hallmarks of aging noninvasively. Here, we demonstrate that cellular DNA methylation (DNAm) profiles from fecal samples provide an accurate and reliable molecular clock in wild capuchin monkeys. Analysis of blood, feces, and urine samples from a closely related species shows that DNAm differentiates between species and different types of biological samples. We further find age-associated differences in DNAm relevant to cellular damage, inflammation, and senescence, consistent with hallmarks conserved across humans and other mammalian species, speaking to the comparative potential. By demonstrating that DNAm can be studied non-invasively in wild animals, our research opens new avenues in the study of modifiers of the pace of aging, and increases potential for cross-population and species comparisons.

## Introduction

Aging is a complex and multifaceted process that impacts health, reproduction, and survival–three key components of Darwinian fitness. Strikingly, not all organisms age the same, resulting in diversity in the manifestation, onset, and pace of age-associated decline among similarly-aged individuals ^1–3^. Since chronological age does not fully explain the variability in age-related trajectories observed among individuals, composite measures–often from molecular readouts–that capture “biological age” have been developed to quantify variation in aging within and across individuals ^4,5^. By far, the most widely-used measures of biological aging are epigenetic clocks, which are composite measures of aging built from genome-wide DNA methylation (DNAm) profiles ^4,6^. Epigenetic clocks have proven to be extremely accurate in predicting chronological age across species (e.g., to within 3.3% error relative to the lifespan ^7^), and the difference between epigenetic and chronological age is associated with age-associated disease risk and mortality across species ^4,8^. Outside of its key role in aging ^9^, DNAm is also of broader interest due to its involvement in shaping phenotypic diversity and plasticity ^10,11^. However, to measure DNAm has heretofore required access to primary tissue samples (e.g., blood), which has limited the ability to study age and environmental effects on aging in many wild and non-model species. As a result, we are missing critical opportunities to understand the factors that pattern aging across the tree of life. Taking epigenetic research out of the lab and into the field would not only expand the taxonomic breadth of aging studies but also provide a unique opportunity to contextualize patterns of human longevity and aging within a broader eco-evolutionary framework ^12–14^.

Non-human primates are long-lived and among the most ecologically and socially diverse orders of mammals. They also have a large and growing pool of genomic resources available, making them a valuable taxonomic group for comparative studies across a range of disciplines from anthropology to biomedicine ^15,16^. Furthermore, given their close evolutionary history with humans, and broadly similar aging ^17,18^, they are a highly translationally relevant system for studying aging heterogeneity ^19–24^. Long-running studies of non-human primates, where molecular, social, and ecological environments are deeply documented, are providing key insights into the ecological and social determinants of health and aging ^19,20,22–25^. Naturalistic and wild systems in particular are demonstrating how changing environmental conditions, whether long-lasting, such as drought, or abrupt, such as hurricanes, alter the links between socio-ecological predictors and the pace of aging ^26,27^. However, the promise of comparative studies will only be fully realized if a wide range of species living in diverse ecological contexts is represented. Since most wild populations cannot be sampled regularly for blood or other tissues, methods to quantify the epigenome using non-invasive samples are desperately needed.

Here, we developed and optimized a protocol for quantifying DNAm in non-invasively collected fecal samples from wild white-faced capuchin monkeys (*Cebus imitator*), which are remarkably long-lived (up to 54 years in captivity, up to 37 years in the wild) for their small body size (3-5 kg) ^28^. We found epigenomic signatures of tissues derived from intestinal epithelium, demonstrating that we were capturing molecular signatures of gut epithelial cells from the host animal. Comparisons of DNAm from feces, urine, and blood from a captive population of a closely related capuchin species (*Sapajus apella*) (Fig. 1) revealed that epigenetic profiles varied by species, sex, and sample source. Using only DNAm measured from fecal samples, we developed a highly accurate epigenetic clock that predicted the chronological age of wild capuchins to within 1.59 years (∼3.5% of the capuchin lifespan)–which is on par with the highly accurate blood-based epigenetic clocks developed in humans. Finally, we found intriguing age-associated differences in methylation levels in homeobox genes and variations at genes involved in developmental processes, cell senescence, and immune responses, findings which are broadly consistent with previous research on aging in humans and other mammals ^7,9,11,29^. Taken together, our approach shows that non-invasive measures are a highly informative source of DNAm data, and thus opens the door for study of epigenetic mechanisms in wild animals with high translational potential.

**Fig. 1.**
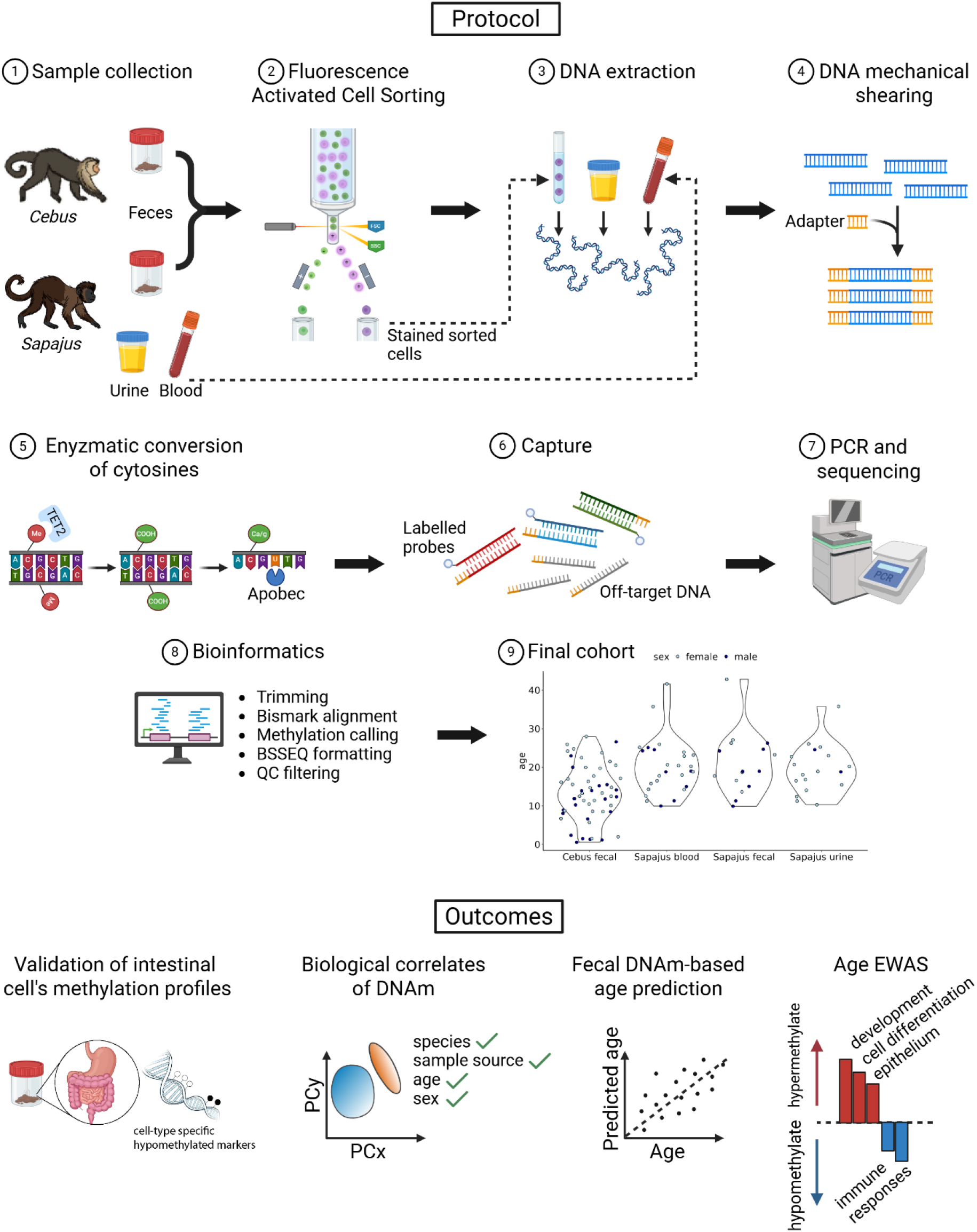
Study design and main outcomes. After collecting fecal samples from wild capuchins (*Cebus imitator*), and fecal, urine, and blood samples from captive capuchins (*Sapajus apella*), we used a flow-cytometry based cell sorting protocol ^30^ to enrich material recovered from fecal samples in capuchin DNA and remove contaminants. We extracted DNA from sorted fecal cells, and from blood and urine samples directly, prior to enzymatic-based methylation sequencing. Following bioinformatics and quality check filtering on libraries, the final cohort included 116 samples across three sample sources and two species. Fecal-derived methylation profiles exhibit signatures of gut tissue specificity, species, age, and sex, and can be leveraged to build accurate methylation clocks. Age-associated differences in methylation from fecal samples hint at conserved variation at a multitude of genomic locations involving housekeeping genes, developmental processes, and immune responses. *Cebus* and *Sapajus* artwork by Jordie Hoffman; organs icons from AdobeStock; created with Biorender.com.

## RESULTS

### Non-invasive sample-derived methylation signatures reflect tissue specificity

We efficiently captured DNA methylation (DNAm) profiles from fecal samples by innovating a Fluorescence-Activated Cell Sorting (fecalFACS) ^30^ plus Twist Targeted Methylation Sequencing (TTMS)–a capture based approach that includes enzymatic methyl sequencing. TTMS uses probes designed to cover ∼4 million CpG sites in the human genome, and we recently demonstrated that this human-based commercial probe set could capture ∼2 million CpG sites in high-quality nonhuman primate samples ^31^ (Table S1). With this novel, combined approach, we covered 905,950 sites in fecal samples from wild-living *Cebus*, and 1,245,571 sites in fecal samples from captive-living *Sapajus* at a coverage of 5x in ≥75% of the samples in the set. This represents about half of the sites recovered from blood samples in this species ^31^, likely due to the more fragmented nature of DNA extracted from fecal samples. After filtering (Table 1), we had high coverage of ∼1 million CpG sites: mean ± SD were 72.21X ± 87.70 in *Sapajus* blood, 184.41X ± 255.53 in *Sapajus* fecal, 161.25X ± 219.63 in *Cebus* fecal, and 316.94X ± 521.37 in *Sapajus* urine samples. For joint analyses across all sample types, we focus on 711,737 CpGs with similar coverage across four datasets.

**Table 1.** Final sample size and demographic characteristics of the cohorts analyzed in this study. For wild *Cebus*, dates of birth were known to within a few days to one month for 81% (n=43 monkeys). The remaining monkeys’ ages were estimated based on morphological characteristics from the first time they were seen.

| Species | Sample source | N° samples | N° individuals | Sex (F/M) | Age (mean±SD) | Age range | Environment |
| --- | --- | --- | --- | --- | --- | --- | --- |
| *Cebus imitator* | fecal | 44 | 43 | 30/13 | 12.19 ± 6.97 | 0.54 - 26.61 | wild |
|  | fecal | 11 | 10 | 3/7 | ~18 ± 6 | ~12 - 28 |  |
| *Sapajus apella* | fecal | 16 | 16 | 8/8 | 21.00 ± 7.96 | 9.90 - 42.84 | captive |
|  | blood | 27 | 27 | 19/8 | 20.30 ± 7.12 | 9.96 - 41.64 |  |
|  | urine | 18 | 17 | 15/2 | 20.00 ± 6.03 | 10.29 - 35.82 |  |

Having effectively captured data from up to 1 million CpG sites in non-invasively collected samples, we next sought to confirm that the DNAm profiles sourced from different biological samples (Table 1) were consistent with the cell types we expected to be present in those samples. To do so we compared our data to a multi-tissue reference DNAm atlas (see Methods), focusing on promoters with cell-specific hypomethylation (table S2). For each sample type, we expected that the tissue of origin (e.g., blood cells in blood-derived samples) would show the strongest hypomethylation in that tissue’s marker genes (e.g., blood marker genes). As expected, blood samples displayed the anticipated low methylation at markers for blood cells (Fig. S1). Fecal-sourced methylation profiles also conformed to expectation. They showed the lowest methylation at marker-specific loci of intestinal epithelium cells (Fig. 2A; 10,000 random permutation p-value = 0.03). We were unable to assess tissue of origin for the urinary samples due to the lack of a urinary tract atlas (Supplemental Results and Fig. S1). Together, these findings demonstrate that: (1) we can effectively quantify DNAm in non-invasively collected samples, and (2) the resulting methylation profiles reflect the biological signatures of the host cells *in situ*.

**Fig. 2.**
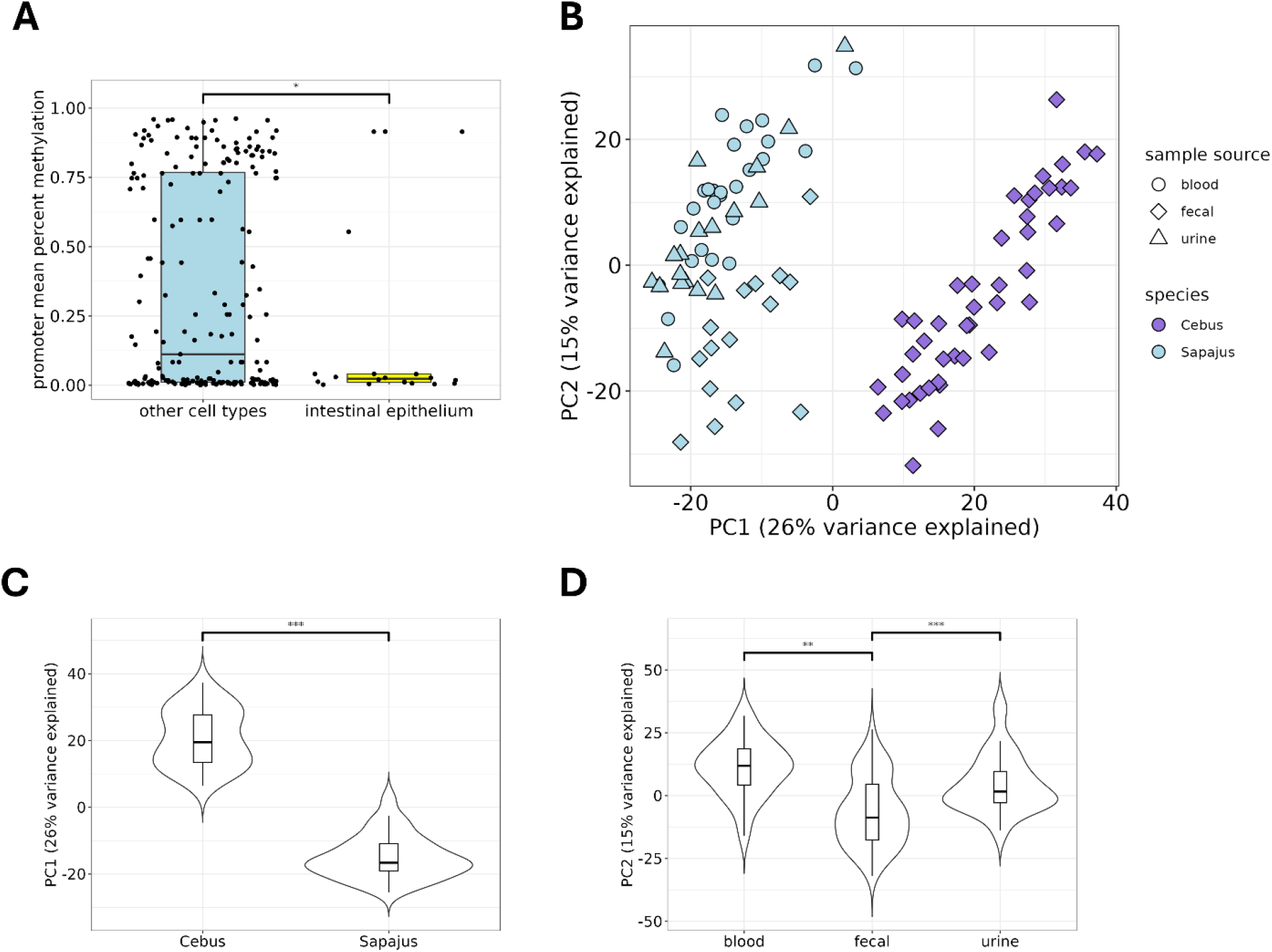
Fecal-derived DNAm profiles capture meaningful biological variation from tissue and species of origin. (**A**) DNAm profiles measured from non-invasively collected fecal samples most closely match the DNAm hypomethylation patterns of intestinal epithelial cells. (**B**) Projection of methylation profiles along the first two components of a PCA. The first principal component significantly partitions samples by species (**C**), while PC2 differentiates fecal from non-fecal samples (**D**). P-values are reported from a linear regression including biological and technical covariates, followed by post hoc pairwise comparisons with correction for multiple testing for factors with more than two levels. In boxplots, boxes represent the interquartile range (IQ), which contains the middle 50% of the records, and a line across the box indicates the median. Vertical lines extend from the upper and lower edges of the box to the highest and lowest values which are no greater than 1.5 times the IQ range. Violin plots display the data distributions and full ranges. P-values are coded as * <0.05, ** <0.01, and *** <0.001.

### DNAm profiles from multiple tissue sources reflect species, sex, and age

We tested whether DNAm profiles would carry meaningful biological signatures, namely of species, age, sex, and sample types. To that end, we used dimensionality reduction by PCA on all samples from individuals for which date of birth was known to within 1 month, thereby excluding 11 samples from 10 *Cebus* (table 1). The first two PCA components (cumulative variance explained = 41%) clearly separated the two species and fecal from non-fecal sample sources (Fig. 2B-D and tables S3-6). Age was included among all top regression models for PC2, demonstrating its important contribution to PC2. Sex was included in 40-50% of the top models for PC1 and PC2 (Fig. S2). These biological differences are unlikely to be due to technical variation: all models controlled for batch or enzymatic conversion effects (tables S3-6 and Fig. S3-4), and analyses included sites with equal mapping rates across the two species to control for reference genome bias (cf. Materials and Methods). Using a multinomial classifier, we could also identify the source of blood, fecal, and urine samples with 96% accuracy (Supplemental results and table S7).

### Accurate age prediction from fecal-derived DNAm profiles

Given the individual-representative biological signatures in the DNAm profiles, we built two DNAm clocks to predict the age of samples based on methylation profiles using elastic net regression. The first clock was trained on all samples from both species (n = 105), maximizing sample size and incorporating data from blood, fecal, and urine samples. The clock was strongly predictive of age, estimating age to within a median of 2.88 years (Pearson’s r=0.82 between chronological and predicted age; Fig. 3A; tables S8-9). Predictions also showed strong reliability within-individuals, with a mean ± SD of 1.65 ± 0.75 years discrepancy across predicted ages for 24 individuals for which we had samples from multiple biological sources. Prediction accuracy was robust to alternative preprocessing steps—such as excluding CpGs with species, sample source, or sex-associated methylation differences–(Methods; Supplemental Results and Fig. S5A). Overall, this performance has comparable accuracy to other DNAm clocks developed in wildlife from blood and other tissues ^7,32,33^.

**Fig. 3.**
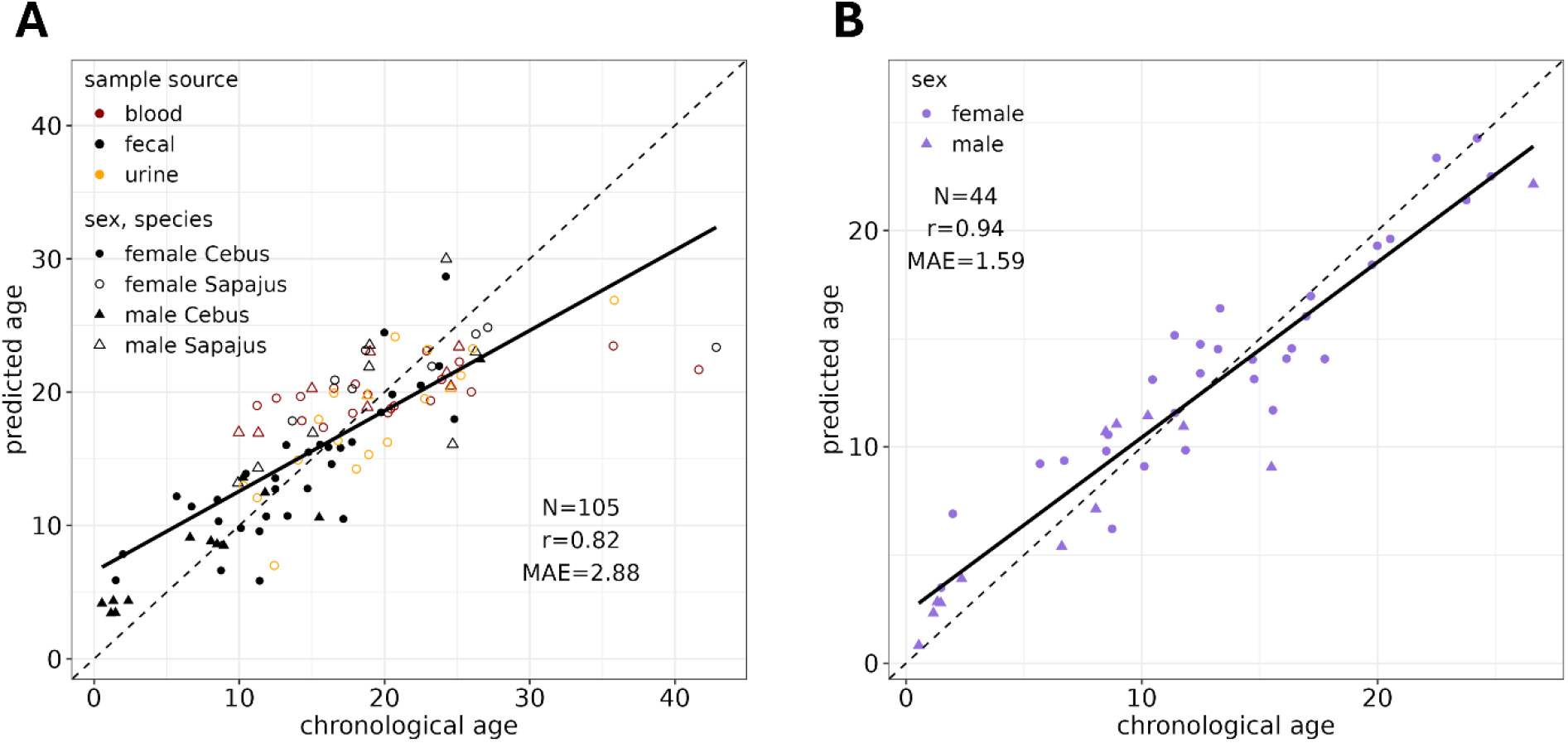
Accurate age estimation from fecal-derived methylation profiles. (**A**) DNAm clock trained on blood, fecal, and urine samples in two capuchin species. Age predictions are from a leave-one-out validation. (**B**) Fecal DNAm clock in *Cebus imitator* estimates the age of wild living individuals from non-invasively collected samples. Model performance is indicated by Pearson’s correlation coefficient (r) and Median Average Error (MAE). The solid lines are the best-fit linear regression of predicted ages on chronological ages, while the dotted lines reflect the line of identity (x=y).

Currently, one of the most pressing challenges for the field is to develop entirely non-invasive techniques for age estimation. To address this challenge, we built a second clock only from fecal samples from the free-ranging white-faced capuchins (*Cebus imitator*). Despite being developed with a smaller sample size, our *Cebus* fecal clock had even higher accuracy than our all-sample clock: with a correlation of 0.94 and median absolute error (MAE) 1.59 years (Fig. 3C; Fig. S5B-6, tables S10-11 and Supplemental Results). When applied to the 11 samples from monkeys with morphologically estimated ages, the clock was less accurate than in monkeys of known age (MAE = 3.5 years; Supplemental Results and Table S12), likely due to imprecision in the morphological estimates. This discrepancy highlights the utility of non-invasive clocks for generating more accurate chronological age estimates, especially in older individuals critical to aging research.

### Fecal-derived methylation profiles captures age-associated differences

While methylation clocks serve as powerful biomarkers by leveraging a small set of CpG sites to predict chronological age, they do not directly illuminate the molecular mechanisms underlying aging. In contrast, identifying CpG sites with age-associated methylation differences can provide insights into gene regulatory changes that accompany aging. Importantly, the ability to generate such data from non-invasively collected samples unlocks new opportunities for comparative aging research across diverse animal species in natural settings.

In service of this goal, we tested for age-associated DNAm differences at 75,521 CpG sites across 60 fecal samples (combined across species). Age was significantly associated with DNAm at 18% of tested sites (n=13,440, FDR < 0.1; table S13). To focus on regions with strong cis-regulatory potential, we mapped 9,754 of these sites to gene promoters (Fig. 4A; Table S14). These age-associated promoters were associated with genes involved in transcriptional regulation, intracellular signaling, immune function, and neural development. For instance, 12 of 14 CpGs in the RPRM promoter—part of the p53 pathway and implicated in gastric cancer risk via methylation-mediated silencing—were significantly hypermethylated ^29,34,35^. Gene ontology analysis further supported these patterns, showing that age-associated hypermethylation was enriched in pathways related to neural function, development, and cell differentiation (Fig. 4B, Fig. S7; Tables S15–17), consistent with previous reports ^7^. In contrast, hypomethylated sites included immune pathways such as TNF and type I interferon signaling, both of which are associated with chronic low-grade inflammation, or “inflammaging” ^36,37^ (Fig. 4B, Fig. S7; Table S15).

**Fig. 4.**
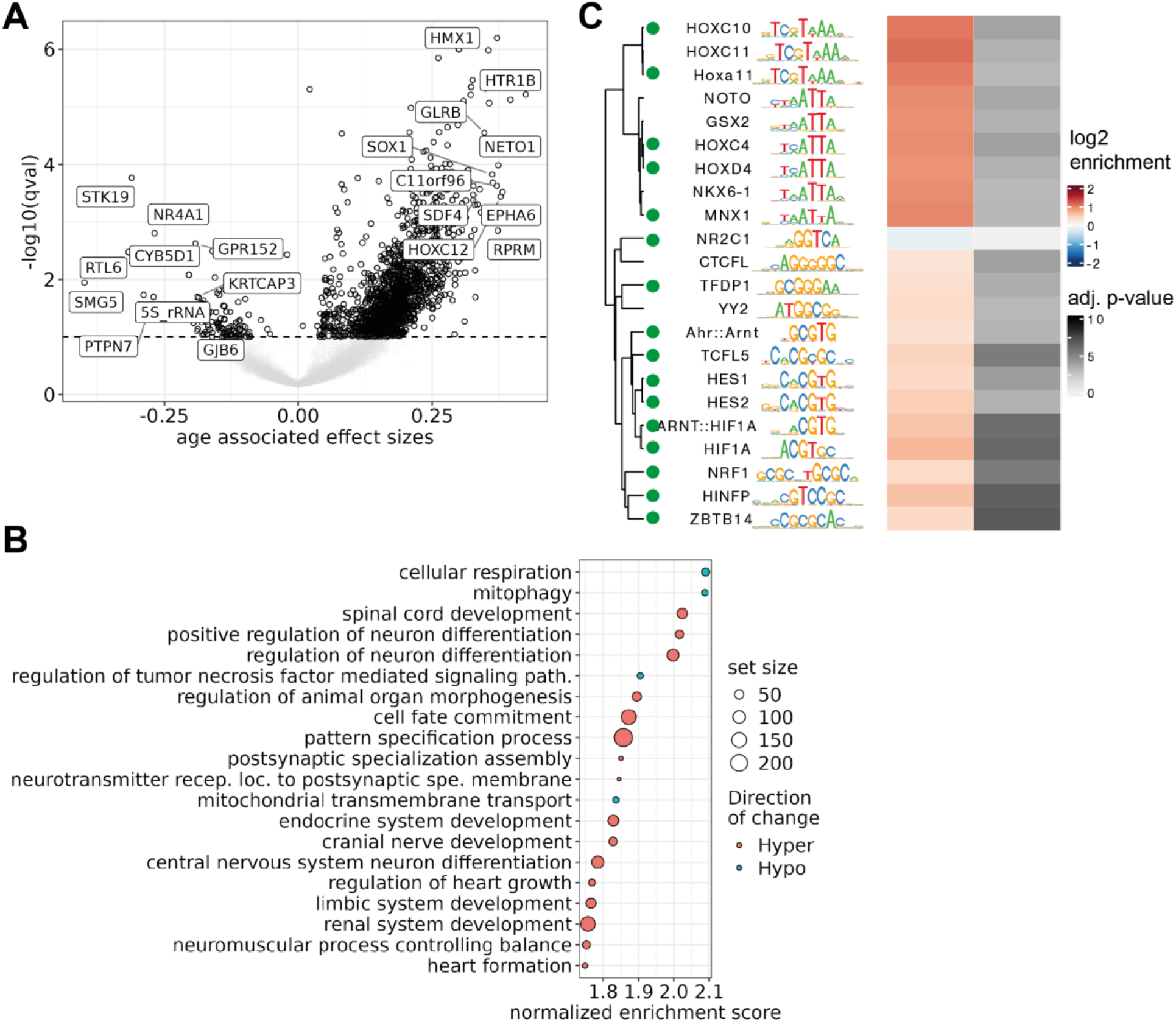
Age is associated with differences in the methylome. (**A**) Age associated differences in methylation levels measured from fecal samples. CpG sites overlapping with promoters in the capuchin genome are represented on the forest plot, with sites reaching FDR <0.1 in black. Top age-associated sites are labelled for illustrative purposes, and include several homeobox genes and protein-coding genes involved in neoplasic processes. (**B**) GO enrichment analysis of the genes overlapping age-associated sites are shown. The 20 most enriched GO terms are shown on the rows, the dot sizes show the number of enriched genes, and the x axis shows the proportion of enriched genes relative to all genes associated with the GO term (absolute normalized enrichment score). All enrichments have Bonferroni-adjusted p-value <0.05. (**C**) Transcription factor binding site enrichment among regions exhibiting higher methylation levels at older ages. Transcription factors expressed in intestinal tissues, based on the Human Protein Atlas, are shown with a green dot.

To investigate potential regulatory impacts, we grouped CpGs into regions and assessed transcription factor binding site (TFBS) enrichment in regions hypermethylated with age. We identified 21 enriched and one under-enriched TFBSs (Fig. 4C), including several involved in development (ZBTB14, HES1, HES2), cell cycle control (TFDP1), and metabolism (NRF1, ARNT::HIF1A). Ten of these TFBSs overlapped with age-associated hypermethylated regions in rhesus macaques ^38^, suggesting some conservation across primates. No enriched TFBSs were found in hypomethylated regions in our dataset.

As an external validation and to assess cross-species conservation, we compared our age-associated CpGs to those identified in a recent multi-tissue pan-mammalian epigenome-wide association study (EWAS) ^7^. Of 3,614 CpGs across 300 genes that could be compared, 69.1% of CpGs that increased in methylation with age in capuchins showed similar directional age-associated differences in the pan-mammalian EWAS, and 73.8% of CpGs that decreased in methylation were similarly consistent (Fig. S8; Table S18). This strong concordance (Fischer’s Exact Test OR = 6.03, 95% Confidence Interval = 5.04 – 7.23, p-value <0.001), despite differences in species and tissue type, demonstrates that reliable insights into methylation can now be drawn for wild species using non-invasive methods.

## Discussion

Understanding the modifiers of aging in wild animals depends on the ability to disentangle biological age from chronological age. Doing so requires access to long-term data from individually monitored populations and biological samples that contain robust aging biomarkers. In this study, we leveraged DNA methylation (DNAm) profiles from cells sorted from non-invasively collected fecal samples in a population of wild capuchin monkeys that has been studied continuously for over 40 years ^28^. We show that chronological age can be estimated with high accuracy—median error of 1.59 years—using fecal DNAm, a substantial improvement over earlier fecal-based efforts in other species, including dolphins ^33^ and mice ^32^, and a microbiome age clock in wild baboons ^39^. Our findings also highlight species-specific and tissue-specific signatures in DNAm, distinguishing not only between two closely related capuchin species but also between feces, blood, and urine samples. These results open new possibilities for non-invasive research— from estimating the ages of unmonitored individuals to exploring how social, environmental, and physiological factors shape the pace of biological aging in different tissues.

Beyond age prediction, we identified thousands of CpG sites with methylation levels that change significantly with age, particularly in gene promoters. These sites were enriched in functional pathways tied to development, neural function, and cell differentiation, consistent with established patterns of epigenetic aging. Conversely, hypomethylated sites with age were enriched in immune-related pathways such as TNF and interferon signaling, implicating processes like inflammation and immune activation—both well-documented hallmarks of aging ^9^. Notably, these patterns were broadly conserved when compared to a recent multi-tissue, pan-mammalian EWAS of age ^7^, reinforcing the comparative value of our findings.

The parallels between our results and those of cross-species EWAS studies support the growing view that aging involves shared molecular signatures, including hypermethylation of genes that regulate cell integrity and transcription, and hypomethylation of immune-related genes. Some of these changes may contribute to processes like senescence or inflammation that are implicated in aging and age-related diseases, including cancer ^40^. The observed enrichment in developmental pathways also aligns with evolutionary theories suggesting that genes beneficial in early life may have detrimental effects later in life ^41,42^. Future work to characterize epigenetic dynamics during development and early life will be key to understanding the origins of variation in aging trajectories.

There is growing evidence that environmental factors shape the pace of aging ^43^, yet we still know relatively little about which factors are the most impactful, which individuals are more susceptible, and which factors may be protective. Long-term studies of wild primates are well-positioned to address this gap, having already linked physical health, hormone levels, social dynamics, and early life adversity to later-life outcomes ^19,20,22,24,28,44,45^. Epigenetic data— especially when derived from blood—has begun to reveal how such exposures leave molecular traces ^26,38^. However, blood is rarely available from wild populations. Our development of non-invasive methods for generating high-resolution DNAm profiles from feces dramatically expands the reach of this approach, enabling comparative and translational studies across species and environments ^33,46,47^.

DNAm is known to reflect both phylogenetic distance and tissue identity ^11,48^. For example, comparisons across humans, chimpanzees, and macaques have shown that methylation profiles cluster strongly by species and organ ^49^. We found that non-invasively collected fecal and urine samples retain such biological signals, demonstrating that field conditions, including sample collection and storage, do not obscure species- and tissue-specific methylation signatures. Standardizing protocols across research teams will be important to further optimize reproducibility and sensitivity under variable field conditions.

We also confirmed that fecal DNAm profiles exhibit canonical hypomethylation at CpG sites specifically associated with epithelial cells of the human lower gastrointestinal tract ^48^. Although DNA from feces and urine likely includes a mix of epithelial cells, immune cells, and cell-free DNA, methylation profiles appear largely preserved ^48,50^. Because DNAm varies by cell type, identifying cellular composition will be essential for accurate biological interpretation. This opens the door to both broad systemic questions and more targeted studies of gut physiology, aging, microbiome interactions, and related metabolic conditions ^25,26,38,51^—many of which are central to modern human disease.

Overall, we have shown that non-invasively collected fecal samples provide a reliable source of DNAm profiles for the study of aging in wild populations. The addition of epigenetic processes to the toolkit available to field research has the potential to bolster our understanding of the modifiers of the pace of aging across a range of environments. The inclusion of less-represented species in aging research with high translational relevance is necessary to maximize the impact of findings related to pace of aging. More broadly, non-invasive epigenetic research will accelerate discovery across domains, from developmental plasticity and resilience to environmental stressors to the evolution of aging in natural contexts.

## Materials and Methods

### Methylation profiles using non-invasive sampling Ethics and Authorization

Fecal samples from wild capuchins in Costa Rica were collected under permits issued by the Animal Care Committee (ACC) of the University of Calgary in Canada (AC19-0167/AC24-0021), and by the Sistema Nacional de Áreas de Conservación (SINAC) and the Área de Conservación Guanacaste (ACG: R-SINAC-ACG-PI-059-2022/ ACG-PI-033-2023ACG-PI-011-2024/, and CONAGEBIO (R-013-2022-OT-CONAGEBIO/R-042-2023-OT-CONAGEBIO) in Costa Rica. Fecal samples were imported to Canada under Canadian Food Inspection Agency (CFIA) permits A-2023-06194-1 and A-2022-05488-4.

Blood, fecal and urine samples were collected from captive brown capuchins (*Sapajus apella*) at Georgia State University, under IACUC protocol (A20018). Fecal and urine samples were imported to the University of Calgary, Canada, under CFIA permit A-2024-03380-4.

### Study populations and sampling

We collected fecal samples from 53 habituated, wild white-faced capuchins (*Cebus imitator*) in Sector Santa Rosa, Área de Conservación Guanacaste, Costa Rica. Capuchins in this population have been studied nearly continuously since the Santa Rosa Primate Project was initiated in 1983 (reviewed in ^28^). All individuals were individually identified through fur patterns, scars, and natural morphological variation. We sampled 20 males and 33 females. The majority of individuals (42/53) have been followed since birth and their date of birth is known to within 1 day - ca. 1 month. One female first seen as an infant had only a known year of birth. We considered ages to be known for these 43 individuals. Ages ranged from 6 months to 26 years old in males, and from 1 year old to 24 years old in females. Other individuals (males N = 7, females N=3) were tracked from the time they were first observed as subadults or adults (for males, typically when they immigrated into the study population; table S1). An experienced researcher estimated the ages of these 10 individuals, and we excluded them from all analyses including age as a predictor.

Approximately 1 g of feces was collected from forest substrates immediately following defecation by trained and experienced researchers wearing nitrile gloves and a face mask, and transferred into a 5 mL conical tube containing 2.5 mL of RNAlater. The samples were stored at room temperature until they were shipped to the University of Calgary for processing.

Brown capuchins (*Sapajus apella*) were members of the captive capuchin monkey colony at Georgia State University. This colony was originally formed in 2006, and currently contains 8 males (ages 11-26 at the time of sample collection) and 20 females (ages 22-42 at the time of sample collection). Monkeys live in mixed sex social groups, excepting one bachelor pair of males, and most monkeys have lived with their group mates their entire lives. Each group, including the bachelor pair, has a dedicated indoor room and outdoor yard, to which they have access except during voluntary testing and inclement weather. Monkeys are fed a species typical diet including monkey chow, fruits, vegetables, and treats, and monkeys have access to water *ad libitum*. Urine and fecal samples were collected from clean trays placed beneath the monkeys’ testing areas during voluntary behavioral and cognitive testing for other research. Samples were placed in 5 mL of RNAlater and stored at room temperature until they were shipped to the University of Calgary for processing. Monkeys are never restricted from food, water, treats, outdoor access, or social contact to motivate participation in research; as a result, urine and fecal samples were only available from monkeys who chose to participate in testing. Whole blood samples were collected during the annual physicals conducted under anesthesia using 13 mg/kg Ketamine, delivered intramuscularly by the veterinarian team. Blood samples were stored at 4ºC upon collection, and shipped to Arizona State University where they were flash frozen into 0.5 mL aliquots and stored at -80 ºC.

### Flow cytometry

We followed our previously validated method for sorting primate cells from the fecal matrix using flow cytometry ^30^, with a few modifications to optimize cell recovery. In brief, we homogenized the fecal samples in RNAlater by vortexing for 30 seconds, then centrifuged at 1,727 rpm for 15 seconds to pellet the larger material. We transferred the supernatant to a 15 mL tube and filled it with Dulbecco’s phosphate-buffered saline (DPBS). We then filtered the supernatant through a 70 μm filter into a 50 mL tube. We transferred the resulting filtrate into a 15 mL tube and centrifuged at 1,500 rpm for 5 minutes to pellet the cells. We washed the pellet twice with 13 mL of DPBS. Then, resuspended the pellet in 300 μL of DPBS and filtered the solution again through a 35 μm filter into a 5 mL FACS tube. We prepared a negative control by mixing 250 μL of DPBS with 50 μL of the cell solution to account for autofluorescence. Next, we added 250 μL of 12 μM DAPI stain and 3 μL of AE1/AE3 Pan cytokeratin Alexa Fluor 488 antibody (ThermoFisher: 53-9003-82). We incubated the samples at room temperature for 15 minutes, followed by an incubation at 4 °C for 15 min to 1 hour, depending on time to initiate flow cytometry.

The cells were isolated using a BD FACSAria Fusion (BD Biosciences) flow cytometer at the University of Calgary Flow Cytometry Core with BD FACSDiva™ Software. Background fluorescence and cellular integrity were assessed by processing the negative control sample before all other prepared fecal samples. For each sample, we first gated the target population based on forward- and side-scatter characteristics to minimize the presence of bacteria and cellular debris. Second, additional secondary and tertiary gates were applied to eliminate cellular agglomerations. Finally, we selected cells with antibody or DNA fluorescence that exceeded background levels. In instances where staining was ineffective, sorting was performed using only the first three gates.

### DNA extraction and quality assessment

Post sorting, we extracted DNA from fecal cells using the Arcturus PicoPure DNA Extraction Kit (Thermo Fisher Scientific, Kit # 0103), according to the manufacturer’s instructions. We extracted DNA directly from urine (*Sapajus* only) stored in RNAlater using the same kit and protocol. Following the extractions, a cleanup step was carried out using Sera-Mag Speedbeads (Fisher Scientific, catalog # 09-981-123) at a 1.5:1 ratio. We extracted DNA from blood samples (*Sapajus*) at Arizona State University using the Qiagen DNeasy Blood & Tissue kits (Qiagen #69581) following the manufacturer’s protocols.

### TMS library preparation and sequencing

We prepared 147 libraries (cf. Table 1 in main text) for TMS library preparation and sequencing. Detailed descriptions of the protocols can be found in ^31^. In brief, we used 200 ng of DNA as the input for library prep with NEBNext Enzymatic Methyl-seq kit (P/N: E7120L). Library prep was modified to eight cycles of PCR for the final library amplification followed by a 0.65X SPRI bead cleanup. Libraries were then combined in equimolar amounts into pools of 12 (total concentration of 2,000 ng per pool) for capture using the Human Methylome panel from Twist Biosciences following the manufacturer’s instructions (P/N: 105521). Post-hybridization libraries were then sequenced on the NovaSeq 6000 at the Vanderbilt Technologies for Advanced Genomics (VANTAGE) Core using 150 bp paired-end sequencing, with a target of 30-50 M paired-end reads per sample.

### Sequence alignment and processing

Paired-end FASTQ files were trimmed using Trim Galore! (function trim_galore with the flag --paired) https://www.bioinformatics.babraham.ac.uk/projects/trim_galore/ and mapped in bismark ^52^ (functions bismark, --score_min L,0,-0.6 -R 10 \ -p 4) to the reference genome for the Panamanian White-faced Capuchin Cebus_imitator-1.0 (https://www.ncbi.nlm.nih.gov/datasets/genome/GCF_001604975.1/). Cytosine methylated and total counts were extracted with functions bismark_methylation_extractor and coverage2cytosine (flag --merge_CpG). Data quality assessments were performed using MultiQC (v1.7; Illumina). We assembled count data using the package *bsseq* (function read.bismark) ^53^ for further processing and analysis in RStudio version 4.4.0 ^54^. We first removed samples with a conversation rate >2% at CHH or CHG (n=1), then those with rate of mapping to the reference genome <50% were excluded from further analysis (n=19 samples: 6 fecal *Cebus*, 6 fecal *Sapajus*, and 8 urine) as well as one library with excessively high sequencing depth. Finally, we removed duplicated libraries (n = 10 excluded). The final sample size was 71 fecal, 27 blood, and 18 urine samples (table 1).

The rate of mapping tended to be lower for *Sapajus* samples relative to *Cebus* samples, which was expected as our reference genome was the *Cebus* genome assembly (Wilcoxon rank sum test W = 2035, p-value = 0.048, fig. S3). To ensure that any differences in the methylomes between species or sample sources would not result from this bias, we filtered CpGs in the four datasets separately (*Cebus*-fecal, *Sapajus*-fecal, *Sapajus*-urine, and *Sapajus*-blood) using a threshold of 5x in ≥75% of the samples in the set. This ensured that all CpGs included in the final data had adequate coverage in every species-sample-source subset. Then, we intersected the subsets for common CpG sites according to the type of data included for each analysis. For example, analyses using blood, feces, and urine used the intersection of the four independently filtered datasets, while analyses using only feces used the intersection of the two fecal sample subsets.

### Methylation signatures of tissue specificity

DNA methylation (DNAm) patterns are highly specific to cell type, such that measurements taken from bulk tissue or blood samples largely reflect the composition of cell types present in those samples. This cell-type specificity poses a challenge when analyzing unconventional media such as fecal and urine samples, where the diversity and representation of cell types are less documented. We anticipated that fecal-sourced host cells would be derived primarily from the intestinal epithelium. However, we wanted to compare the methylation profiles measured from fecal samples to DNAm single-cell profiles to test this assumption and validate our methods. We extracted the top 1000 cell-specific markers identified by the HumanMethylationAtlas ^48^ for all cell types. We focused on the cell-specific hypomethylated loci, because they represent the vast majority of markers identified by Loyfer and colleagues ^48^. Accordingly, a cell-specific marker is a genomic region (one or more sites) which exhibit markedly lower methylation levels in the cell type compared to all other cell types.

Because of the lack of chromosome-level assembly for the capuchin genome, but availability of curated information on genes (gene transfer format or gtf), we focused on annotated gene promoters. We used Cebus_imitator.Cebus_imitator-1.0.113.gtf to extract the location of genes annotated in both the human atlas and the capuchin genome. We passed the bed file coordinate of promoters to the function getCoverage() in *bsseq* with type=“Cov” and “M” with what=“perRegionTotal”, to calculate the coverage and methylation counts over entire promoters. A total of 269 combinations of cell type-promoters from the HumanMethylationAtlas could be associated with 191 promoter sequences from the capuchin genome (table S2). This implies that some promoters were inevitably associated with several cell types (here, promoters were on average considered as hypomethylated markers for mean ± sd = 1.31 ± 0.64 cell types). We calculated promoter mean percent methylation across all samples and compared the mean percent methylation of promoters according to their reference cell-specificity in the HumanMethylationAtlas. We calculated the average difference in percent methylation at promoters annotated as markers for intestinal epithelia versus all other cell types, and used 10,000 permutations of which promoters were assigned as markers of epithelia to create a random distribution of difference in percent methylation between a set of markers and the background of all other cell types. The one-tailed p-value was calculated by comparing the observed and randomized difference.

To further validate our approach, we repeated this procedure with blood samples by investigating the top markers associated with circulatory immune cells and other cell types available from the HumanMethylationAtlas. In total, 231 cell type-promoters from the human data could be matched to 165 promoters in capuchins (1.32 ± 0.63 promoter to cell types).

### Biological variables recovered from multidimensional analysis of methylation profiles

We investigated correlates of methylation profiles using dimensionality reduction with Principal Component Analysis (PCA). We performed PCA using prcomp(scale = TRUE) on the matrix of percent methylation (n = 116 samples) using sites covered across all samples (n = 1,421). To investigate the possible influence of technical artifacts on the biological signal, we visualized the correlations between PC1 and PC2 with sample average percent methylation, conversion rate at CHH, conversation rate at CHG, mapping efficiency, and batch. To assess the relative explanatory power of the biological and technical covariates, we built linear regression models for PC1 and PC2. The Pearson correlation between CHG and CHH methylation (a proxy for enzymatic conversion efficiency) was 0.97, so we only retained CHG, which exhibited greater variance, to avoid issues with multicollinearity. Variance Inflation Factors further revealed that batch number could not be included in a model also including other technical covariates. Therefore, we created two versions of the models: one with batch (as factor), and the other with mapping efficiency, average percent methylation, and conversation rate at CHG. Species, sample source, sex, and age were included in both models. Covariates were z-transformed and models fitted with lm(). We note that some VIFs remained high for species, batch, and sample source, suggesting that fully disentangling the relative influence of these three parameters remains challenging. We then performed AICc-based model comparison using the MuMIn ^55^ dredge(rank = “AICc”) function. We report the proportion of models in which predictors were included among all models within ΔAICc = 10 from the best model. To summarize the outcomes, species was included in all competing best models for PC1, and sample source was included in all the competing best models for PC2, which supports the main findings from the PCA visualization. Loadings of batches on PC1 and PC2 are presented in fig. S4 (tables S3-6).

### Multinomial classifier of sample source

To examine further the discriminatory power of sample source (blood, feces, urine) on methylation profiles, we used a multinomial regression algorithm in *glmnet* ^56,57^ with a leave-one-out validation. The model was trained using cv.glmnet() on all samples but one by mapping sample source against methylation profiles at a set of 106,099 CpG sites with coverage across the four species-sample-source subsets after imputation with missMDA ^58^. The penalization parameter lambda was internally determined by 10 cross-fold validation, and we used predict(type = “response”) with lambda.min on the test sample. The test sample is assigned probabilities that it originates for one sample source or the other. As covariates are not included in *glmnet*, this effectively tests the ability to determine sample source despite noise in the methylation profiles associated with uncorrected sex, age, or technical batch effects.

### Age clocks built from fecal DNA methylation profiles

We used an elastic net regression in *glmnet* with a leave-one-out-validation (LOOV) procedure to achieve the least biased possible estimation of chronological ages based on methylation profiles. A first set of models were fitted on all samples (four species-sample-source combinations, n = 105) to assess model performance on a set of heterogeneous sample sources while maximizing sample size. A second set of models were fitted on fecal samples from the wild *Cebus* of known age (n = 44) to evaluate performance on a smaller sample size of homogeneous samples that speak to our goal of developing non-invasive epigenetic clocks. For both scenarios, we started by imputing missing values using *missMDA* ^58^ (function imputePCA with scale = TRUE and ncp = 2 as determined by estim_ncpPCA with scale = TRUE, ncp.min = 0, ncp.max = 5, method.cv = “Kfold”). Imputation was done independently in each subset, after excluding low variance sites constitutively hypo (average <0.1) or hypermethylated (average > 0.9). From there on, we varied the data preparation process by: i) transforming or not age before sexual maturity, here 5 years old (models Classic and AgeTransfo), ii) normalizing data or not with a Yeo-Johnson transformation, in combination or not with the age transformation (models Norm and AgeTransfoNorm), iii) pre-selecting sites correlating with age >0.2, before normalization and age transformation (models Corr and CorrAgeTransfoNorm), and iv) pre-selecting for sites exhibiting no significant difference in methylation levels according to species and sample sources as found from binomial mixed models (model NoBiasAgeTransfoNorm, see details below) for the clock using all sample sources.

For all LOOV runs, the best penalization parameter lambda was determined internally using 10 cross-fold validation, and we ran iterations across values of alpha between 0 and 1 (i.e., spanning the space from ridge to lasso regression) and chose the alpha minimizing mean squared error (MSE). Model performance is reported based on Pearson’s correlation coefficient and Median Average Error (MAE) expressed in years. For 24 individuals repeatedly sampled (one female *Cebus*, 15 female and eight male *Sapajus*) giving a total of 57 samples (22 blood, 18 fecal, 17 urine), we calculated the level of consistency across predicted ages based on the generated epigenetic clock by calculating for each individual the standard deviation of the absolute error between chronological age and predicted age. We report the mean and standard deviation of this within-individual standard deviation across the 24 individuals.

### Differential methylation between species, sample source, and sex

We tested for loci differential methylation using MACAU implemented in *PQLseq* ^59–61^. Our aim was to identify and remove loci (N = 69,353) which may lower clock’s performance due to confounding effects of species, sample source, or sex on methylation levels. We ran binomial mixed models on data generated from fecal samples to test for a difference between *Cebus* and *Sapajus*, while accounting for dummy-coded sex and age (models converged for N = 42,939 CpGs). Among samples collected from *Sapajus*, we tested for a difference in blood versus fecal and urine, while accounting for dummy-coded sex and age (models converged for N = 58,697 CpGs). Finally, we tested for an effect of sex among fecal samples collected in *Cebus*, while accounting for age (models converged for N = 52,623 CpGs). All models included an identity matrix as a relatedness matrix, with relatedness set to 1 for samples from the same individual. For each set of results and after inspecting the distribution of p-values, we calculated q-values using the package *qvalue* ^62^ which corrects for the number of tests performed. We did not add batch effects because our aim here is to test for an effect of species on data as raw as possible which is the format passed on to *glmnet*. From all three procedures, we extracted 20,313 sites which exhibited q-values > 0.05 in all cases (i.e., not statistically influenced by sex, sample source, or species) for the NoBiasAgeTransfoNorm elastic net regression model.

### Age-associated differences in fecal methylation profiles

To test for loci differential methylation with age, we followed a similar procedure to the one described for species, sample source, and sex. Models estimating the effect of age on fecal methylation profiles included dummy-coded sex, species, and batch as covariates, and an identity matrix. Models that did not converge (59%) were excluded from downstream analysis, leaving 75,521 loci (table S10). Here again q-values were calculated from p-values ^62^. To identify putative age-associated differences in gene expression with age, we more closely examined a subset of 9,754 sites which overlapped with promoters from the capuchin gtf annotation (table S10). The genomic location for the promoters were extracted from the capuchin’s gtf file using makeTxDbFromGFF() and genes() in the package *GenomicFeatures*, and promoters(upstream=2000, downstream=200) from *IRanges*. We intersected these genomic coordinates with results from our analysis on age-associated changes in capuchin (function findOverlap() with default settings). To allow comparisons with previous panmammalian studies, we also extracted the meta-analysis effect sizes for age at CpG overlapping gene promoters from Lu et al. ^7^ and annotated our results at overlapping promoters.

### Pathway enrichment

To test for enrichment of molecular and cellular pathways, we focused on n = 52,887 CpGs overlapping genes. The effect sizes were averaged across CpGs overlapping the same promoters, leaving 5,239 genes which we ranked by effect sizes. We performed pathway enrichment analyses using fgsea() from the eponym package ^63,64^ (arguments minSize = 15, maxSize = 500, nPermSimple = 10000, eps = 0.0), with the subcollection Gene Ontology Biological Processes, Cellular Components, and Molecular Functions retrieved from msigdbr(species = “Homo sapiens”) ^65,66^. For plotting, we simplified the results using collapsePathways() on pathways reaching an FDR <0.05 and show the top 20 pathways ranked by absolute normalized enrichment scores.

### Transcription factors enrichment

To be able to test for the presence of DNA motifs known to be associated with the binding of transcription factors, we first grouped single CpGs into differentially methylated regions (DMRs) using the pipeline from ^38,51^ available at (https://github.com/mwatowich/Immune-gene-regulation-is-associated-with-age-and-environmental-adversity-in-a-nonhuman-primate). DMRs were defined as segments including a higher-than-expected density of significantly age-associated sites exhibiting consistent direction of change. First, we determined the number of CpG sites loosely associated with age (FDR<0.1) within 1,000 base pairs of an age-associated site (FDR<0.05) at chance level by randomly permuting p-values among CpGs (median of three loosely age-associated CpGs per DMR). Therefore, we kept from the real data DMRs with at least four loosely age-associated sites. Then, DMRs were removed if fewer than 75% of the CpGs in the DMR or significant CpGs at the loose threshold were in the same direction. We filtered nine DMRs which were longer than 99% of the distribution, leaving 876 DMRs, which were 475 ± 433 base pairs long on average (range = nine – 2,149), and included 18 ± 15 CpGs (range = four – 117), of which 11 ± 8 and 8 ± 7 were loosely and strictly age-associated respectively. We defined a background set of DMRs by applying no threshold on significance and concordance of age-associated changes, and further removing the set of previously identified DMRs from the background set. Finally, we filtered background regions longer than the longest DMR (n=87 regions removed) to obtain more homogeneous sets (4,374 background regions of average length 455 ± 420, range = 10 – 2,230).

We tested for transcription factors binding site motifs (TFBSs) enrichments using *monaLisa* ^67^. We downloaded vertebrates’ transcription factor binding site position weight matrices from Jaspar 2020 ^68^. Region sequences were extracted from the capuchin genome converted to a FAFile using getSeq() from *BSgenome* ^69^. We compared hypermethylated DMRs (n = 844) and hypomethylated DMRs (n = 32) to the background set. TFBSs with adjusted p-value < 0.001 are shown.

TFs expression is tissue-dependent, which implies that enrichment for TFBSs motifs at differentially methylated sites in the cells from the intestinal tract are likely to have consequences for TFs expressed in that tissue. Using the Human Protein Atlas ^70^ (https://www.proteinatlas.org/), we considered that a TFs was likely to be expressed in the gastrointestinal tract if the protein expression score was medium or high or if the RNA expression consensus listed digestive tissues among the top third of the tissues.

### Overlap with age-associated sites in pan mammalian EWAS of age

We extracted the top 1000 CpGs exhibiting higher and 1000 CpGs exhibiting lower methylation with age in an EWAS across several tissues and species of eutherian mammals from Lu and colleagues (2023) ^7^. We reasoned that these top age-associated sites are more likely to be shared broadly. Because we could not map CpGs from the pan mammalian array to the capuchin genome directly, we relied on gene annotations. Specifically, we used gene names from the pan mammalian data to extract the genomic location for the promoters and exons present in the capuchin’s gtf file (package *IRanges* functions promoter() and exonsBy() with by = gene). We intersected these genomic coordinates with results from our analysis on age-associated changes in capuchin (function findOverlap() with default settings), which returned 3,772 CpGs overlapping with a gene annotation. We did not apply any filtering for significance as we are interested in the consistency of the direction of difference broadly. Also, note that some genes could be associated with several CpGs, which may or may not differ in the same direction with age. To compare the direction of difference in the capuchin and pan mammalian dataset, we had to ensure that each gene would be represented by a unique direction of difference with age in the pan mammalian data. Genes that were unambiguously associated with effect sizes all in one direction were first selected (n = 786). Then, for genes associated with several CpGs changing in opposite directions (n = 65), we calculated the proportion of sites that are higher with age. After visual inspection of the distribution, we decided to keep sites exhibiting a proportion of positive differences <25% and >75% (n = 30). Sites falling between these boundaries were excluded. We assigned the direction of difference of the majority of CpGs overlapping the gene. This procedure allowed us to assign either a positive or negative direction of difference with age for 816 genes out of the 851 originally present in the pan mammalian dataset. We then intersected the datasets to compare the direction of difference for 300 genes common to both datasets (n = 3,614 CpGs, with an average 12.0 ± 14.6 CpGs overlapping a gene in capuchins) (table S18).

## Authors’ contributions

Conceptualization: Melin, Snyder-Mackler

Methodology: Melin, Snyder-Mackler, Hernández-Rojas, Hamou, Orkin, Sadoughi

Validation: Melin, Snyder-Mackler, Brosnan, Sadoughi

Formal analysis: Sadoughi, Hernández-Rojas, Hamou

Investigation: Melin, Sadoughi, Snyder-Mackler, Hernández-Rojas, Hamou,

Resources: Melin, Jack, Campos, Brosnan, Higham, Snyder-Mackler

Data collection: Lopez, Simmons, Mah, Slikas

Data curation: Campos, Melin, Hernández-Rojas, Hamou, Mah, Snyder-Mackler, Sadoughi

Writing – original draft: Sadoughi, Melin, Snyder-Mackler

Writing – review & editing: all authors

Visualization: Sadoughi, Melin

Supervision: Melin, Campos, Jack, Snyder-Mackler

Project administration: Melin, Campos, Jack, Snyder-Mackler, Mah

Funding acquisition: Melin, Campos, Jack, Higham, Orkin, Snyder-Mackle

All authors read and approved the manuscript.

## Data and materials availability

Genomic sequences generated as part of this study have been deposited in NCBI’s Sequence Read Archive and will be accessible upon publication. R code and bash command lines are available from github https://github.com/BaptisteSadoughi/CapuchinsDNAm.

## Acknowledgements

We would like to thank Roger Blanco, Maria Marta, Staff and Administration of ACG at SSR. Linda M Fedigan, Saul Cheves Hernandez, Danielka Rugama Taylor, Wendy Tellez Arias, Philippine Delga, Lindy Wolhuter, Cielo De La Rosa Meza, Anda Pohle, Peyton Schidli, Suheidy Romero Morales, and all members of the Santa Rosa field team. Patricia Ströher for administration support. The GSU veterinarian, Dr. Rex Howard, and other members of the Department of Animal resources for their assistance collecting blood samples of the *Sapajus* and Kelly Leverett for coordination and support. Christine Adjangba, Sarah van Dijk, and Ashlee Greenier for lab support, and members of the Snyder-Mackler lab for constructive feedback on earlier versions of the manuscript.

This work was supported by funding from the Natural Sciences and Engineering Research Council of Canada (NSERC; RGPIN-2023-04399, and DGECR-2023-00272 to JDO, RGPIN-2017-03782 to ADM, CGS to MEM), the Canada Research Chairs program (950-231257 to ADM), the Natiohnal Institutes of Health Aging Research In Animals (NIH ARIA) program (R61-AG078529-01 to FC, ADM, KJ, JPH, JDO, NSM), and the National Science Foundation (NSF BCS 2127375 and NSF SES 1919305 to SFB).

## Notes

### Competing Interest Statement

The authors have declared no competing interest.

## References

1. Avila-Rieger, J. et al. Socioeconomic Status, Biological Aging, and Memory in a Diverse National Sample of Older US Men and Women. Neurology 99, e2114–e2124 (2022).

2. Fiorito, G. et al. Social adversity and epigenetic aging: a multi-cohort study on socioeconomic differences in peripheral blood DNA methylation. Sci. Rep. 7, 16266 (2017).

3. Kivimäki, M. et al. Social disadvantage accelerates aging. Nat. Med. 1–9 (2025) doi:10.1038/s41591-025-03563-4.

4. Horvath, S. & Raj, K. DNA methylation-based biomarkers and the epigenetic clock theory of ageing. Nat. Rev. Genet. 19, 371–384 (2018).

5. Rutledge, J., Oh, H. & Wyss-Coray, T. Measuring biological age using omics data. Nat. Rev. Genet. 23, 715–727 (2022).

6. Bell, C. G. et al. DNA methylation aging clocks: challenges and recommendations. Genome Biol. 20, 249 (2019).

7. Lu, A. T. et al. Universal DNA methylation age across mammalian tissues. Nat. Aging 3, 1144–1166 (2023).

8. Levine, M. E. et al. An epigenetic biomarker of aging for lifespan and healthspan. Aging 10, 573–591 (2018).

9. López-Otín, C., Blasco, M. A., Partridge, L., Serrano, M. & Kroemer, G. Hallmarks of aging: An expanding universe. Cell 186, 243–278 (2023).

10. Greenberg, M. V. C. & Bourc’his, D. The diverse roles of DNA methylation in mammalian development and disease. Nat. Rev. Mol. Cell Biol. 20, 590–607 (2019).

11. Haghani, A. et al. DNA methylation networks underlying mammalian traits. Science 381, eabq5693 (2023).

12. Newediuk, L. et al. Designing epigenetic clocks for wildlife research. (2025).

13. Nussey, D. H. Studying aging in the wild can help us to understand resilience and healthy aging. Nat. Aging 5, 337–338 (2025).

14. Roach, D. A. & Carey, J. R. Population Biology of Aging in the Wild. Annu. Rev. Ecol. Evol. Syst. 45, 421–443 (2014).

15. Fiziev, P. P. et al. Rare penetrant mutations confer severe risk of common diseases. Science 380, eabo1131 (2023).

16. L Rocha, J., Lou, R. N. & Sudmant, P. H. Structural variation in humans and our primate kin in the era of telomere-to-telomere genomes and pangenomics. Curr. Opin. Genet. Dev. 87, 102233 (2024).

17. Bronikowski, A. M. et al. Aging in the Natural World: Comparative Data Reveal Similar Mortality Patterns Across Primates. Science 331, 1325–1328 (2011).

18. Colchero, F. et al. The long lives of primates and the ‘invariant rate of ageing’ hypothesis. Nat. Commun. 12, 3666 (2021).

19. Campos, F. A. et al. Wild capuchin monkeys as a model system for investigating the social and ecological determinants of ageing. Philos. Trans. R. Soc. B Biol. Sci. 379, 20230482 (2024).

20. Emery Thompson, M., Muller, M. N., Machanda, Z. P., Otali, E. & Wrangham, R. W. The Kibale Chimpanzee Project: Over thirty years of research, conservation, and change. Biol. Conserv. 252, 108857 (2020).

21. Fernandes, A. G., Poirier, A. C., Veilleux, C. C. & Melin, A. D. Contributions and future potential of animal models for geroscience research on sensory systems. GeroScience 47, 61–83 (2025).

22. Newman, L. E. et al. The biology of aging in a social world: Insights from free-ranging rhesus macaques. Neurosci. Biobehav. Rev. 154, 105424 (2023).

23. Tung, J., Lange, E. C., Alberts, S. C. & Archie, E. A. Social and early life determinants of survival from cradle to grave: A case study in wild baboons. Neurosci. Biobehav. Rev. 152, 105282 (2023).

24. Wilson, M. L. et al. Research and conservation in the greater Gombe ecosystem: challenges and opportunities. Biol. Conserv. 252, 108853 (2020).

25. Sadoughi, B., Schneider, D., Daniel, R., Schülke, O. & Ostner, J. Aging gut microbiota of wild macaques are equally diverse, less stable, but progressively personalized. Microbiome 10, 95 (2022).

26. Anderson, J. A. et al. DNA methylation signatures of early-life adversity are exposure-dependent in wild baboons. Proc. Natl. Acad. Sci. 121, e2309469121 (2024).

27. Testard, C. et al. Ecological disturbance alters the adaptive benefits of social ties. Science 384, 1330–1335 (2024).

28. Melin, A. D. et al. Primate life history, social dynamics, ecology, and conservation: Contributions from long-term research in Área de Conservación Guanacaste, Costa Rica. Biotropica 52, 1041–1064 (2020).

29. Wang, H. et al. Methylation-Sensitive Melt Curve Analysis of the Reprimo Gene Methylation in Gastric Cancer. PLOS ONE 11, e0168635 (2016).

30. Orkin, J. D. et al. The genomics of ecological flexibility, large brains, and long lives in capuchin monkeys revealed with fecalFACS. Proc. Natl. Acad. Sci. 118, e2010632118 (2021).

31. Longtin, A. et al. Cost-effective solutions for high-throughput enzymatic DNA methylation sequencing. PLOS Genet. 21, e1011667 (2025).

32. Hanski, E. et al. Epigenetic age estimation of wild mice using faecal samples. Mol. Ecol. 33, e17330 (2024).

33. Yagi, G. et al. Non-invasive age estimation based on faecal DNA using methylation-sensitive high-resolution melting for Indo-Pacific bottlenose dolphins. Mol. Ecol. Resour. 24, e13906 (2024).

34. Liu, L. & Yang, X. Implication of Reprimo and hMLH1 gene methylation in early diagnosis of gastric carcinoma. Int. J. Clin. Exp. Pathol. 8, 14977–14982 (2015).

35. Wu, Y. & and Du, J. Downregulated Reprimo by LINC00467 participates in the growth and metastasis of gastric cancer. Bioengineered 13, 11893–11906 (2022).

36. De Cecco, M. et al. L1 drives IFN in senescent cells and promotes age-associated inflammation. Nature 566, 73–78 (2019).

37. Page, M. J., Kell, D. B. & Pretorius, E. The Role of Lipopolysaccharide-Induced Cell Signalling in Chronic Inflammation. Chronic Stress 6, 24705470221076390 (2022).

38. Watowich, M. M. et al. Immune gene regulation is associated with age and environmental adversity in a nonhuman primate. Mol. Ecol. 33, e17445 (2024).

39. Dasari, M. R. et al. Social and environmental predictors of gut microbiome age in wild baboons. eLife 13, (2024).

40. Vincze, O. et al. Cancer risk across mammals. Nature 601, 263–267 (2022).

41. de Magalhães, J. P. Ageing as a software design flaw. Genome Biol. 24, 51 (2023).

42. Lemaître, J.-F. et al. Early-late life trade-offs and the evolution of ageing in the wild. Proc R Soc B 282, 20150209 (2015).

43. Argentieri, M. A. et al. Integrating the environmental and genetic architectures of aging and mortality. Nat. Med. 1–10 (2025) doi:10.1038/s41591-024-03483-9.

44. Boesch, C., Boesch, P. C. & Boesch-Achermann, H. The Chimpanzees of the Taï Forest: Behavioural Ecology and Evolution. (Oxford University Press, 2000).

45. Perry, S., Godoy, I. & Lammers, W. The Lomas Barbudal Monkey Project: Two Decades of Research on Cebus capucinus. in Long-Term Field Studies of Primates (eds. Kappeler, P.M. & Watts, D.P.) 141–163 (Springer Berlin Heidelberg, Berlin, Heidelberg, 2012). doi:10.1007/978-3-642-22514-7_7.

46. Nick Weber, D. et al. Noninvasive, epigenetic age estimation in an elasmobranch, the cownose ray (Rhinoptera bonasus). Sci. Rep. 14, 26261 (2024).

47. Vullioud, C. et al. Epigenetic signatures of social status in wild female spotted hyenas (Crocuta crocuta). Commun. Biol. 7, 1–12 (2024).

48. Loyfer, N. et al. A DNA methylation atlas of normal human cell types. Nature 613, 355–364 (2023).

49. Blake, L. E. et al. A comparison of gene expression and DNA methylation patterns across tissues and species. Genome Res. 30, 250–262 (2020).

50. Banila, C. et al. A noninvasive method for whole-genome skin methylome profiling. Br. J. Dermatol. 189, 750–759 (2023).

51. Lea, A. J., Altmann, J., Alberts, S. C. & Tung, J. Resource base influences genome-wide DNA methylation levels in wild baboons (Papio cynocephalus). Mol. Ecol. 25, 1681–1696 (2016).

52. Krueger, F. & Andrews, S. R. Bismark: a flexible aligner and methylation caller for Bisulfite-Seq applications. Bioinformatics 27, 1571–1572 (2011).

53. Hansen, K. D., Langmead, B. & Irizarry, R. A. BSmooth: from whole genome bisulfite sequencing reads to differentially methylated regions. Genome Biol. 13, R83 (2012).

54. RStudio Team. RStudio: Integrated Development for R. RStudio, PBC, Boston, MA. (2022).

55. Bartoń, K. MuMIn: Multi-Model Inference. (2024).

56. Friedman, J. H., Hastie, T. & Tibshirani, R. Regularization Paths for Generalized Linear Models via Coordinate Descent. J. Stat. Softw. 33, 1–22 (2010).

57. Tay, J. K., Narasimhan, B. & Hastie, T. Elastic Net Regularization Paths for All Generalized Linear Models. J. Stat. Softw. 106, 1–31 (2023).

58. Josse, J. & Husson, F. missMDA: A Package for Handling Missing Values in Multivariate Data Analysis. J. Stat. Softw. 70, 1–31 (2016).

59. Lea, A. J., Vilgalys, T. P., Durst, P. A. P. & Tung, J. Maximizing ecological and evolutionary insight in bisulfite sequencing data sets. Nat. Ecol. Evol. 1, 1074–1083 (2017).

60. Lea, A. J., Tung, J. & Zhou, X. A Flexible, Efficient Binomial Mixed Model for Identifying Differential DNA Methylation in Bisulfite Sequencing Data. PLOS Genet. 11, e1005650 (2015).

61. Sun, S., Zhu, J. & Zhou, X. PQLseq: Efficient Mixed Model Analysis of Count Data in Large-Scale Genomic Sequencing Studies. 1.2.1 (2021).

62. Storey, J., Bass, A., Dabney, A. & Robinson. qvalue: Q-value estimation for false discovery rate control. (2024).

63. Korotkevich, G. et al. Fast gene set enrichment analysis. 060012 Preprint at 10.1101/060012 (2021).

64. Subramanian, A. et al. Gene set enrichment analysis: A knowledge-based approach for interpreting genome-wide expression profiles. Proc. Natl. Acad. Sci. 102, 15545–15550 (2005).

65. Liberzon, A. et al. The Molecular Signatures Database Hallmark Gene Set Collection. Cell Syst. 1, 417–425 (2015).

66. Liberzon, A. et al. Molecular signatures database (MSigDB) 3.0. Bioinformatics 27, 1739– 1740 (2011).

67. Machlab, D. et al. monaLisa: an R/Bioconductor package for identifying regulatory motifs. Bioinformatics 38, 2624–2625 (2022).

68. Fornes, O. et al. JASPAR 2020: update of the open-access database of transcription factor binding profiles. Nucleic Acids Res. 48, D87–D92 (2020).

69. Pagès, H. BSgenome: Software infrastructure for efficient representation of full genomes and their SNPs. doi:10.18129/B9.bioc.BSgenome, R package version 1.76.0,. (2025).

70. Thul, P. J. et al. A subcellular map of the human proteome. Science 356, eaal3321 (2017).

