## Supplemental information and figures for "Non-invasive measures of DNA methylation capture molecular aging in wild capuchin monkeys"

### Supplemental results

#### Methylation signatures of tissue specificity

The HumanMethylationAltas includes bladder epithelium as a category. However, few hypomethylated markers were identified, and none overlapped with gene promoters present in our capuchin methylation profiles. Therefore, we could not map urine samples to a directly analogous tissue type from HumanMethylationAtlas. Rather, we considered markers for epithelial cells from any tissues as a potential target. However, methylation levels at these promoters were not hypomethylated (Fig. S1). This was expected because epithelial cells from different body locations exhibit different methylation marks in humans [48]. The main limitation of our approach lies in the necessity to map the promoters of analogous genes, instead of lifting over from humans to capuchins the precise CpG coordinates. A refined chromosome-level assembly and annotation of the capuchin genome will allow to more precisely map cell-type markers and explore the specificity of cells recovered from urine samples in the future.

#### Multinomial classifier of sample source

To examine further the discriminatory power of sample source (blood, feces, urine) on methylation profiles, we used a multinomial regression algorithm in *glmnet* with a leave-one-out validation. For each sample, the model assigns a probability that it originated from one of the three sources. The model correctly assigned samples in 96% of the cases (with one fecal and four urine samples that were misclassified). Along with findings reported in the main text, these results confirm that strong biological signals are recovered from DNA methylation (DNAm) profiles measured not only from blood, but also from fecal and urine samples.

#### Comparison of all samples and *Cebus* fecal age clocks' performance

We compared the performance of several versions of the clock trained on all sample sources in both species, and of several versions of the clock trained on fecal samples in *Cebus,* by means of Pearson's correlation coefficient with age, and MAE. As explained in the Material and Methods, the different clock versions differed in the steps involved in the data pre-processing. For the clock trained on all samples, we found that the best clock included normalization of methylation levels and transformation of age with respect to age at sexual maturity (Fig. S5A). For *Cebus* fecal samples, the best clock only involved a pre-selection step of CpGs based on Pearson's correlation between percent methylation and age (Fig. S5B).

The clock trained on *Cebus* fecal samples outperformed the clock trained on all sample sources, despite a larger sample size in the latter. We wanted to rule out that this lower performance in the clock trained on all sample sources could be due to the inclusion of a few very old individuals that are poorly predicted (cf. Fig. 3A). Therefore, we reran the best version of the clock after excluding samples from individuals above 27 years old, thereby matching the age range across the clocks. The performance of the clock trained on all samples increased significantly (Fig. S6) but remained below that of the *Cebus* fecal clock. Therefore, the gap in performance is more likely explained by the greater homogeneity among methylation profiles recovered from one sample source in one species compared to three sample sources and two species.

**Ages as estimated by an experienced observer and based on DNA methylation profiles**

The 11 ages estimated by an observer and those derived from the *Cebus* fecal clock showed strong agreement overall (MAE = 3.5 years). Based on the year of first observation, it is possible to identify cases where the clock is likely to be more, or less, accurate than the observer (table S12). For five individuals, the two estimations are within 3 years and can be considered accurate. One female and two males have a discrepancy between 3.5 and 7 years, but both estimated ages are compatible with their morphological aspect at first sight; true ages are most likely somewhere within the boundaries. For the remaining two females and one male, the clock estimation is very unlikely or incompatible with the individual physical appearance at first sight (with one individual predicted to be a very young juvenile while it was estimated to be about five, and two predicted to have been juveniles or very young adults while they were described as already aged individuals). This is not surprising knowing that these three individuals are at the upper end of the age range used to train the clock, and that predictions are accurate when the training and test sets encompass the similar age ranges [12].

**Supplemental figures**

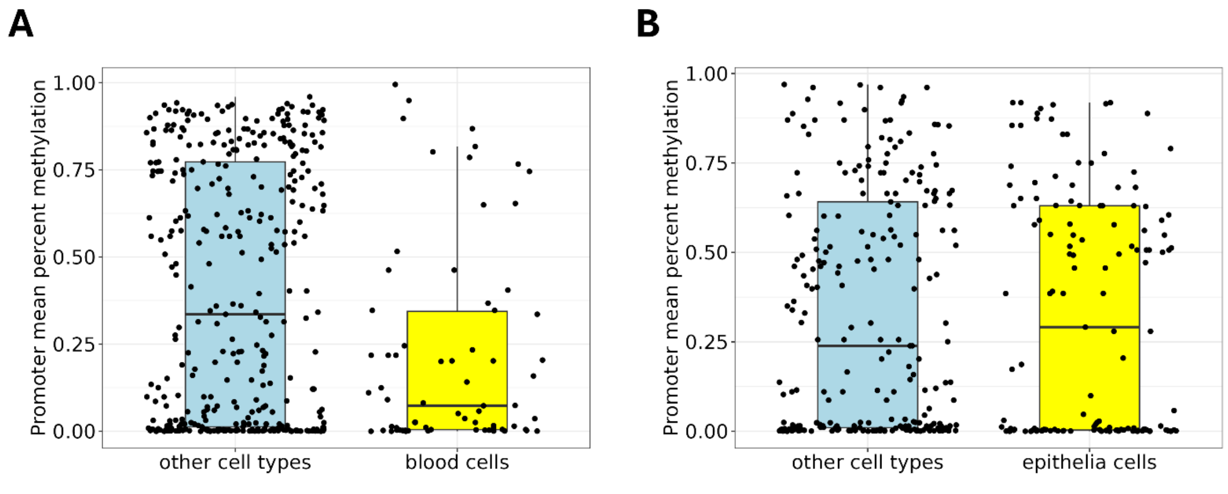


**Fig. S1. Percent methylation at cell-specific hypomethylated markers.** (**A**) In blood and (**B**) in urine samples. Cell-specific hypomethylated markers were defined based on the HumanMethylationAtlas. Boxes represent the interquartile range (IQ), which contains the middle 50% of the records, and a line across the box indicates the median. Vertical lines extend from the upper and lower edges of the box to the highest and lowest values which are no greater than 1.5 times the IQ range.

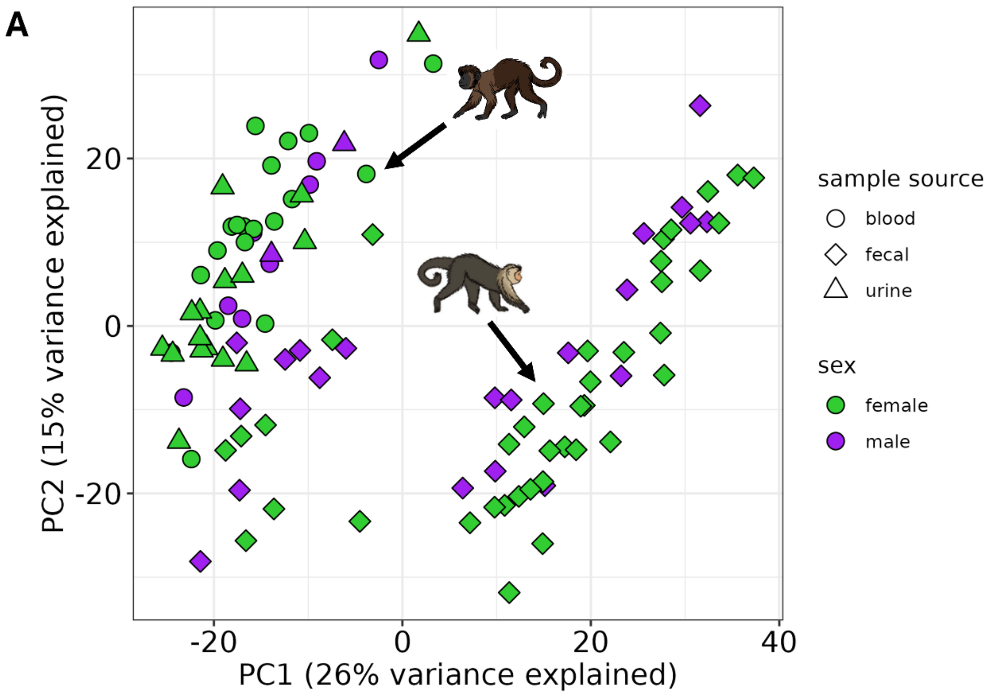


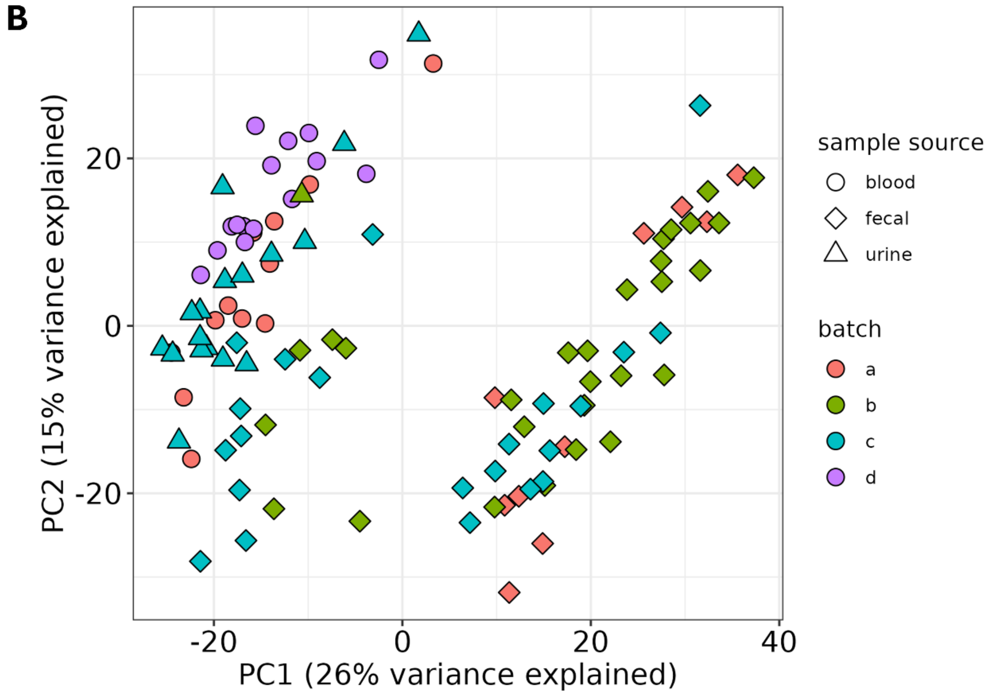


**Fig. S2. Projection of methylation profiles along the first two components of a PCA.** Samples are colored by sex (**A**) or batch (**B**).

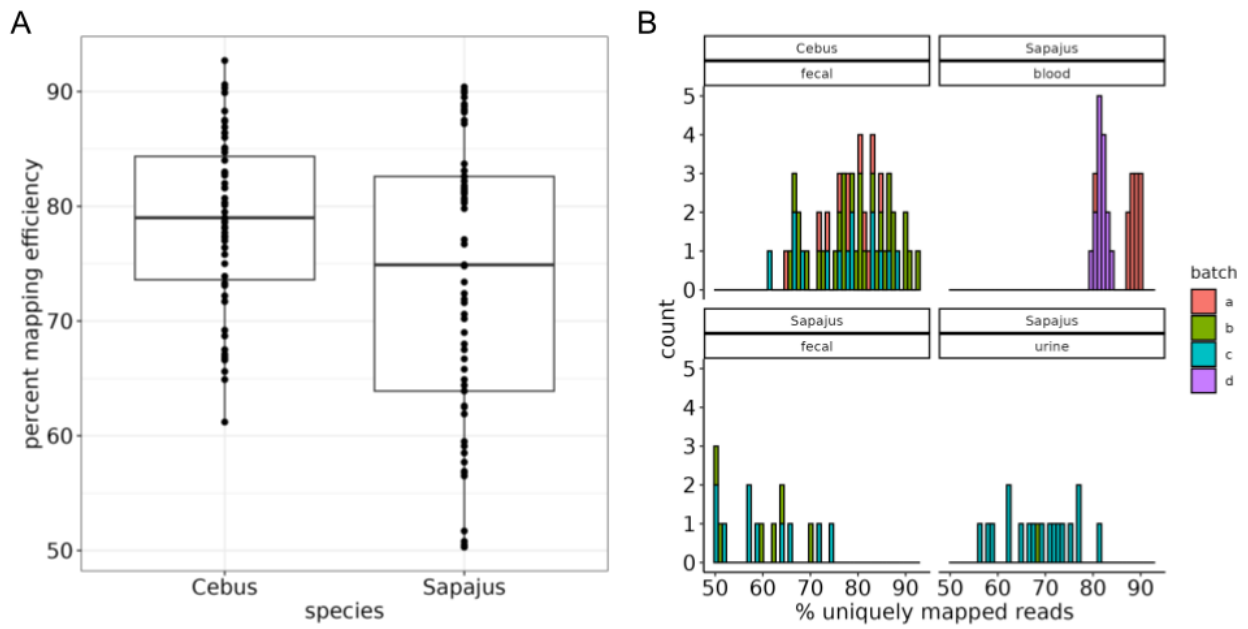


**Fig. S3. Mapping efficiency in Cebus and Sapajus across batches.** (**A**) Mapping was lower in Sapajus, which could be partially explained by the fact that the reference genome used for alignment was from Cebus. Boxes represent the interquartile range (IQ), which contains the middle 50% of the records, and a line across the box indicates the median. Vertical lines extend from the upper and lower edges of the box to the highest and lowest values which are no greater than 1.5 times the IQ range. (**B**) Batch effects coupled with uneven representation of each species across batches might also contribute to the apparent species differences.

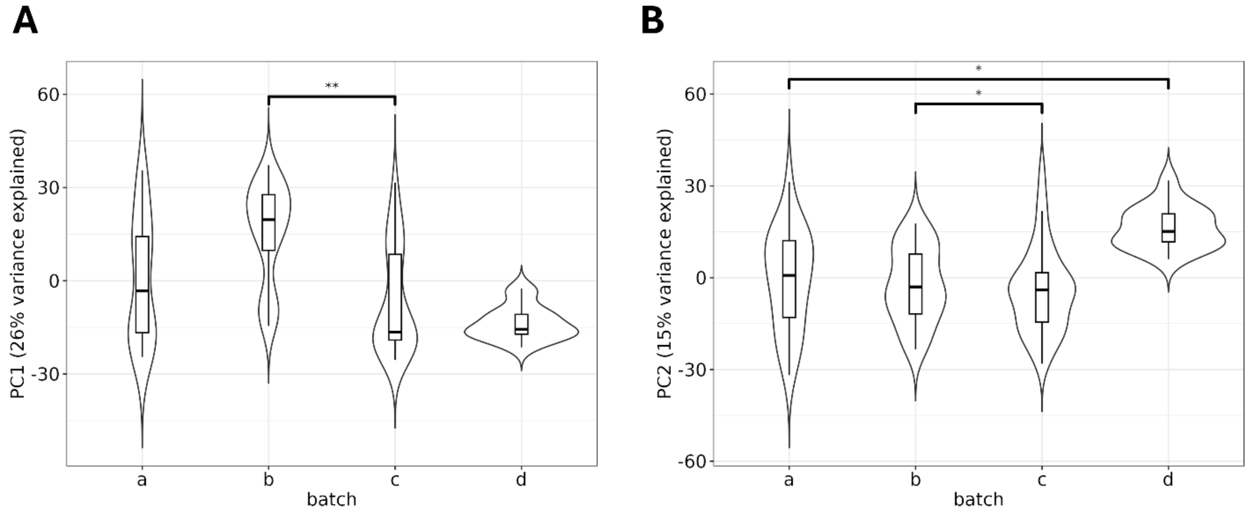


**Fig. S4. Batch effects on methylation profiles.** Boxplots and violin plots of values among four batches are plotted against **(A)** PC1 and **(B)** PC2. The effect of batch on PC1 and PC2 loading was assessed using model comparison which showed that batch was included in the best competing models. To extract p-values for visual representation, we ran a linear model including all predictors, and ran post hoc pairwise comparisons on adjusted marginal means in package *emmeans* with Tukey correction for multiple testing (n=4). Boxes represent the interquartile range (IQ), which contains the middle 50% of the records, and a line across the box indicates the median. Vertical lines extend from the upper and lower edges of the box to the highest and lowest values which are no greater than 1.5 times the IQ range. Violin plots display the data distributions and full ranges. P-values are coded as * <0.05, ** <0.01, and *** <0.001.

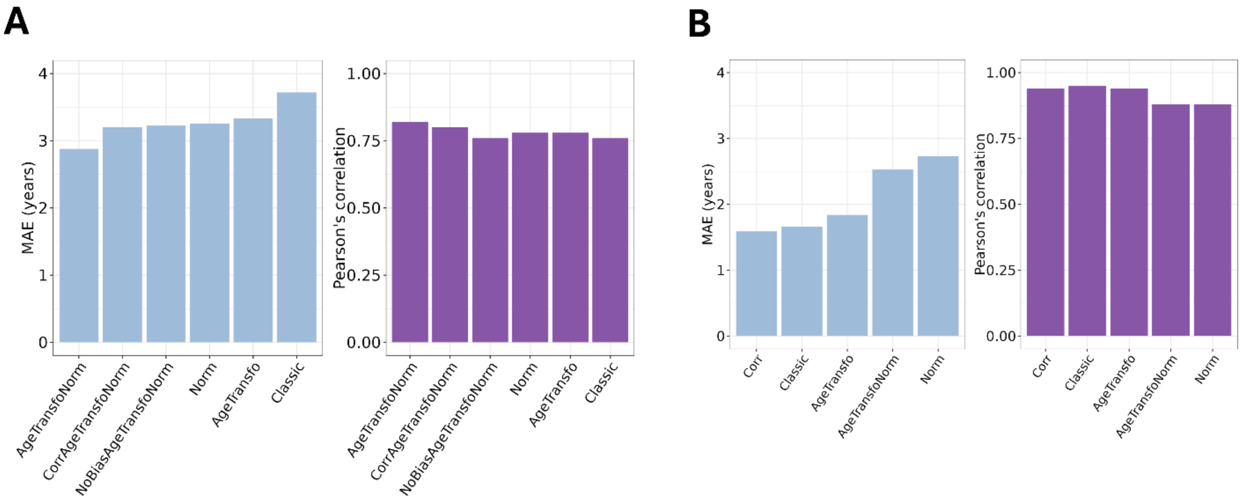


**Fig. S5. Performance of methylation clocks.** Elastic net regression algorithms were trained on (**A**) all sample types, and (**B**) *Cebus* fecal samples only with different levels of pre-processing. Models are ordered on the x-axis per increasing Median Average Error (MAE) expressed in years.

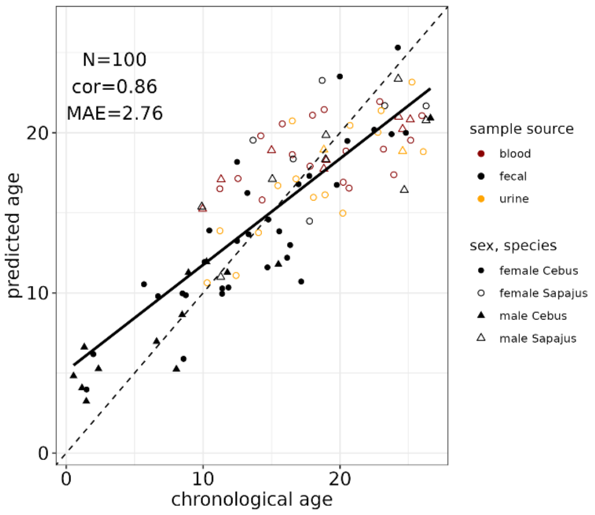


**Fig. S6. Improvements of the clock trained on all samples.** The best performing version of the clock trained on all sample sources was rerun after excluding the oldest individuals in order to match age ranges between clock types. Model performance is indicated by Pearson's correlation coefficient (cor) and MAE, with N the sample size. The full lines show a perfect match between chronological and predicted ages, and the dotted lines the linear regression of predicted ages on chronological ages.

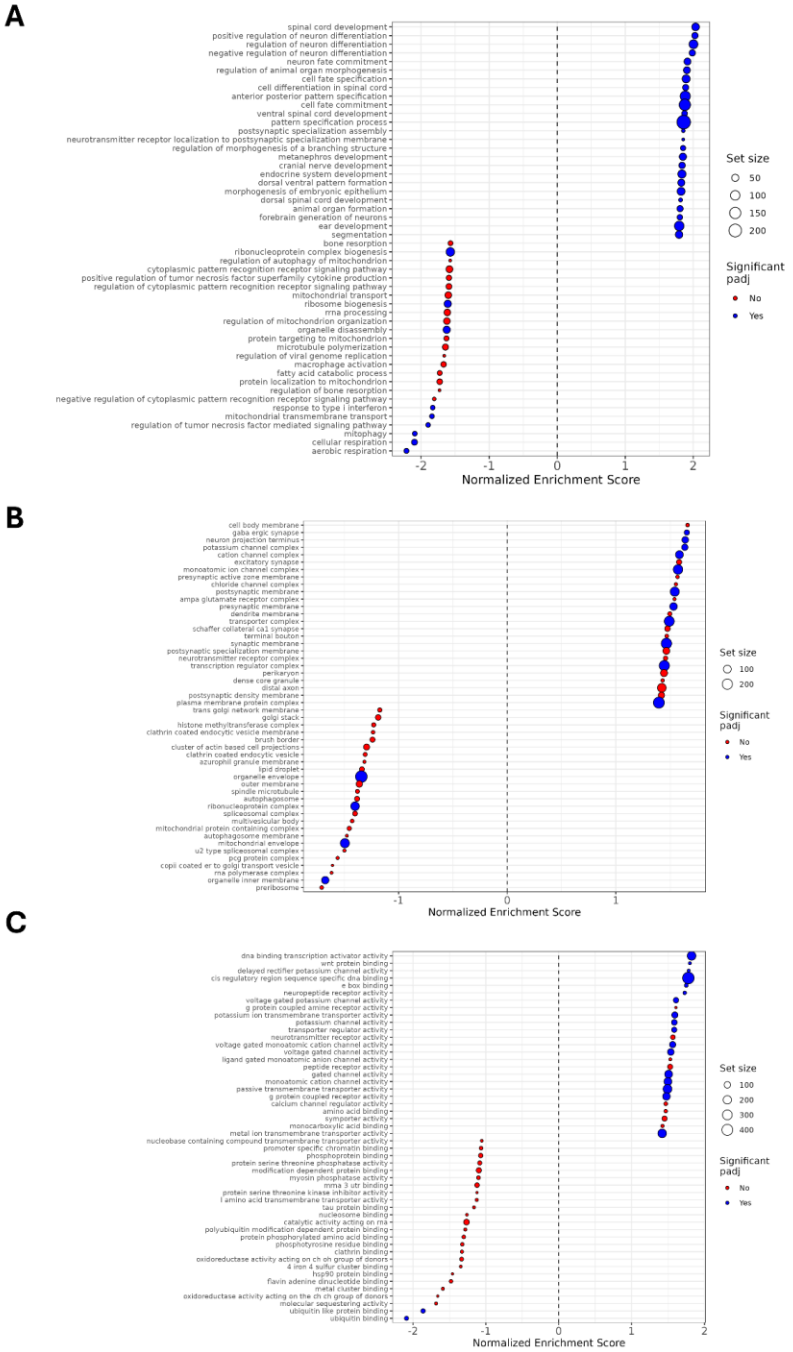


**Fig. S7. Pathways enrichment with age.** Top age-associated enriched pathways for (**A**) biological processes, (**B**) molecular functions, and (**C**) cellular components. The top 25 positively and 25 negatively enriched pathways are plotted.

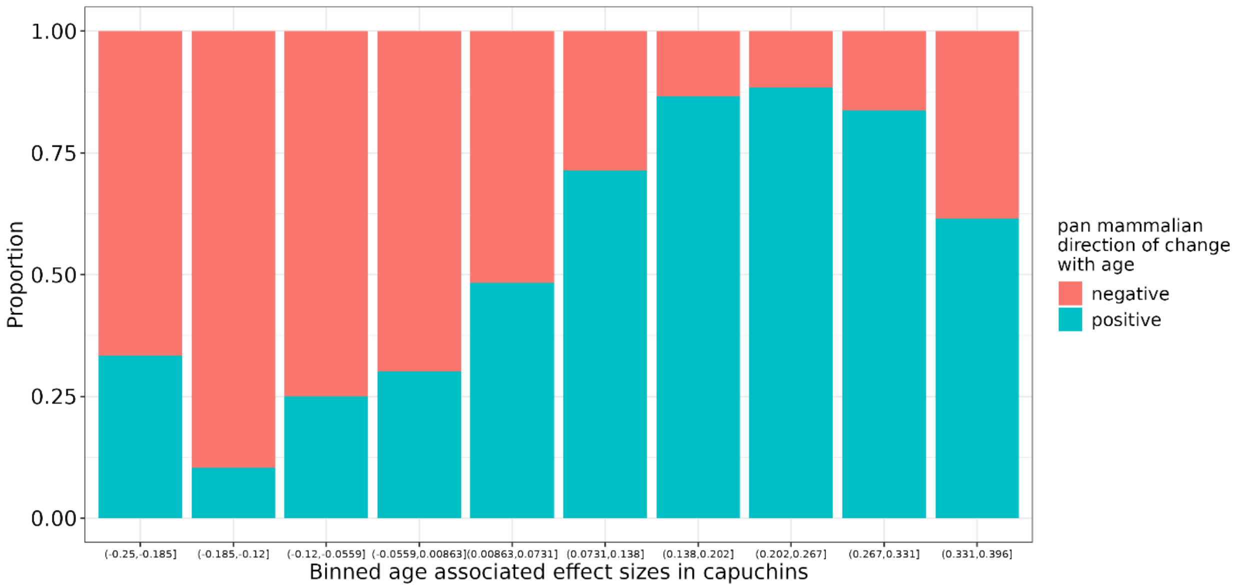


**Fig. S8. Consistency of age-associated changes in capuchin's fecal methylomes and pan tissue samples from mammals.** CpG sites were included if they overlapped a gene or promoter which was also covered in the mammalian dataset. The x-axis shows binned effect sizes for the effect of age in capuchins. The y-axis shows the corresponding direction of change with age in the pan mammalian analysis of age [7].

### Supplementary Files

Tables S1-18 in excel format.
